# Proliferative response of ERα-positive breast cancer cells to 10 μM enterolactone, and the associated alteration in the transcriptomic landscape

**DOI:** 10.1101/2023.04.16.537055

**Authors:** Juana Hatwik, Hrishikesh Nitin Patil, Anil Mukund Limaye

**Author notes:** Corresponding author Anil Mukund Limaye; Department of Biosciences and Bioengineering Indian Institute of Technology Guwahati Guwahati 781039, Assam, India.

## Abstract

Enterolactone (EL) is a product of gut-microbial metabolism of dietary plant lignans. Studies linking EL with breast cancer risk have bolstered investigations into its effects on the mammary epithelial cells, and the mechanisms thereof. While it binds to the estrogen receptor α; ERα, its effect on the proliferation of mammary tumor cell lines is reportedly ambivalent; depending on its concentration. The genomic correlates of EL actions also remain unexplored. Here we have elaborately studied the effect of EL on proliferation of ERα-positive, and ERα-negative cell lines. 10 µM EL significantly enhanced the growth of the ERα-positive MCF-7 or T47D breast cancer cells, but not the ERα-negative MDA-MB-231 or MDA-MB-453 cells. In MCF-7 cells, it significantly increased the expression of *TFF1* mRNA, an estrogen-induced transcript. The binding of ERα to the estrogen response element within the *TFF1* locus further demonstrates the pro-estrogenic effect of 10 µM EL. We further explored the genome-wide transcriptomic effect of 10 µM EL. Analysis of RNA-seq data obtained from control- or 10 µM EL treated-MCF-7 cells revealed modulation of expression of diverse sets of functionally related genes, which reflected cell cycle progression, rather than cell cycle arrest or apoptosis. The manner in which 10 µM EL regulated the hallmark G2/M checkpoint, and estrogen-response-late genes correlated with proliferation inducing, and estrogen-like effects of EL on MCF-7 cells.

## 1. Introduction

Enterolactone (EL), a mammalian enterolignan is a product of gut-microbial transformation of dietary plant-lignan precursors (Borriello et al., 1985; Setchell et al., 1981). It is derived directly from matairesinol, or indirectly from secoisolariciresinol, via the formation of enterodiol (Adlercreutz, 2002). EL, which is detectable in serum of individuals on routine diet (Adlercreutz et al., 1994), is significantly elevated in those maintaining vegetarian diets rich in plant lignans, such as flaxseed and sesame (Tarpila et al., 2002). However, the absolute level of serum EL is determined by gut microflora, along with other factors such as constipation, and intake of coffee, tea, and alcohol (Horner et al., 2002; Kilkkinen et al., 2001).

EL is believed to be a key mediator of health promoting effects of plant lignan-rich diets; including breast cancer risk reduction (Touré and Xueming, 2010). However, the literature is divided on the association of plant lignan intake, or serum EL concentration, with breast cancer risk. Independent epidemiological studies have produced conflicting results; either in favour of, or against any association (Boccardo et al., 2004; Kilkkinen et al., 2004; Pietinen et al., 2001; Zeleniuch-Jacquotte et al., 2004). The causal relationship between EL exposure and breast cancer risk is still in the realm of hypothesis, and bereft of mechanistic insights. Case-control studies on the relationship between serum EL and breast cancer risk, have yielded mixed results (Pietinen et al., 2001; Zeleniuch-Jacquotte et al., 2004). More recent systematic reviews, or meta-analyses have revealed limited evidence for the risk-reducing effect of dietary lignans, or serum EL, in post-menopausal, but not in pre-menopausal women (Buck et al., 2010; Liu et al., 2021; Seibold et al., 2014). It is therefore likely that the impact of EL on breast cancer risk may be governed by circulating hormones, or the status of the estrogen-ERα signalling axis in the breast epithelium (Chang et al., 2019; Olsen et al., 2004)

EL binds ERα *in vitro* (Penttinen et al., 2007). It modulates ERα function; producing both estrogenic or antiestrogenic effects (Carreau et al., 2008; Pianjing et al., 2011). These effects were interpreted based on its ability to positively or negatively affect proliferation (Bowers et al., 2019; Mousavi and Adlercreutz, 1992) or induce ERα-target gene expression in ERα-positive breast cancer cells over a wide range of concentration (Pianjing et al., 2011; Sathyamoorthy et al., 1994). Estrogen responsive, and estrogen response elements (EREs) containing reporter assays have revealed pro-estrogenic effects of EL (Carreau et al., 2008; Penttinen et al., 2007). Although the duality in the effect of EL on breast cancer cells is as intriguing as the perceived effects on breast cancer risk, it is likely that both are a result of its interaction with the estrogen-ERα signalling axis (Zhu et al., 2017). The true effect of EL on the breast epithelial cells may be concentration-dependent. Large deviations in serum EL in women on diverse diet and lifestyle regimen (Kilkkinen et al., 2001) may obfuscate its true relationship with breast cancer risk.

Despite the availability of the next generation sequencing technology, the genome-wide transcriptomic correlates of EL action, particularly with reference to its interaction with ERα signalling, or its proliferation inducing effects in breast cancer cells, have remained unexplored. This limits our understanding of EL’s true impact on breast cancer etiology. Here we show that 10 µM EL induces proliferation of ERα-positive breast cancer cells, which is blocked by tamoxifen. Taking a cue from the 10 µM EL-mediated induction of *TFF1* mRNA expression in MCF-7 breast cancer cells, which is associated with ERα binding to the estrogen responsive element in the *TFF1* locus, we employed next generation sequencing to capture the transcriptomic response of MCF-7 cells treated with 10 µM EL.

## 2. Materials and Methods

### 2.1. Plasticwares, chemicals, and reagents

Plasticwares were purchased from Eppendorf (Hamburg, Germany) and Thermo Fisher Scientific (PA, USA). Roswell Park Memorial Institute Medium (RPMI-1640), Dulbecco Modified Eagle Medium (DMEM), Dulbecco’s phosphate-buffered saline (DPBS), Fetal bovine serum (FBS), charcoal-stripped fetal bovine serum (CS-FBS), Trypsin-EDTA, and antibiotics were from HiMedia Laboratories (Mumbai, India). Enterolactone (EL, Cat. No. 45199), and 17β-estradiol (E2, Cat. No. E8875) were purchased from Sigma-Aldrich (St. Louis, MO, USA). All other reagents, salts, and buffers were purchased from Merck (Darmstadt, Germany), Sisco Research Laboratories Pvt Ltd (Mumbai, India) and Sigma-Aldrich (St. Louis, MO, USA). EL was dissolved in dimethylsulfoxide (DMSO) to a working stock concentration of 80 mM was prepared, and stored at −20°C.

### 2.2. Cell lines and culture media

MCF-7, T47D, MDA-MB-231, and MDA-MB-453 breast cancer cell lines were procured from the National Centre for Cell Sciences, Pune, India. For routine expansion, the cells were grown in DMEM (for MCF-7) or RPMI-1640 (for T47D, MDA-MB-231, and MDA-MB-453) supplemented with 10% FBS, 100 units/mL penicillin, and 100 μg/mL streptomycin (M1 medium). For experiments involving treatment with E2 or EL, we used phenol red-free DMEM or RPMI-1640, supplemented with 10% CS-FBS, 100 units/mL penicillin, and 100 μg/mL streptomycin (M2 medium).

### 2.3. Cell viability assays

Cell lines were seeded in 35 mm dishes using M1 medium at a density of 7×10^3^ cells/dish. After 48 h, the cells were fed M2 medium for 24 h, before commencing the experiments. For dose-response studies, cells were fed with M2 medium supplemented with varying concentrations of EL (0 to 40 µM) or 10 nM E2 for 0 h (baseline viable cell count) or 120 h (end-point viable cell count) in separate sets of culture dishes. Cells treated for 120 h were fed with fresh treatment medium every 24 h. For time-course studies, the cells were treated with vehicle (0.1% DMSO), 10 nM E2 or 10 µM EL in M2 medium for 0, 24, 48, 72, 96 and 120 h in separate sets of dishes. Upon completion of the experiments, the cells were washed with DPBS, trypsinized, and suspended in 1 mL medium. The cell suspensions were mixed with equal volume of 0.4% trypan blue dye (Sigma-Aldrich, St. Louis, MO, USA, Cat No. T8154), and viable cells were counted using a hemocytometer.

### 2.4. Western blotting

Total protein was extracted in 1.5X Laemmli buffer (Laemmli, 1970), and quantified using the trichloroacetic acid (TCA) method (Karlsson et al., 1994). 30 μg of total protein samples were resolved by 10% denaturing SDS-PAGE, and transferred onto nitrocellulose membranes (HiMedia Laboratories,Mumbai, India, Cat No. SF110B). The membranes were blocked with 1% (w/v) gelatin in Tris-buffered saline containing 0.05% v/v Tween 20 (TBST) for 2 h at room temperature. The blots were probed overnight with anti-PCNA antibody (Cat. No. 13110, Cell Signaling Technology, MA, USA) at 4°C, or for 1 h with anti-H3 antibody (Cat. No. BB-AB0055, BioBharati LifeScience, India) at room temperature. The blots were washed with 1X TBST (6 washes of 5 min each) followed by incubation with HRP-conjugated anti-rabbit secondary antibody (Cat. No. 7074S, Cell Signaling Technology, MA, USA) for 1 h at room temperature. The blots were again washed with 1X TBST (6 washes of 5 min each), and developed by Clarity Western ECL Substrate (Bio-Rad Laboratories, Hercules, CA, USA). The chemiluminescence images were captured with ChemiDoc™ XRS+ System with Image Lab™ Software (Bio-Rad Laboratories, Hercules, CA, USA).

### 2.5. RNA extraction, cDNA synthesis and qPCR

Total RNA was extracted using a reagent prepared in-house as per Chomczynski and Sacchi (Chomczynski and Sacchi, 2006), and quantified using BioSpectrometer basic (Eppendorf, Hamburg, Germany). The integrity of total RNA was checked by agarose gel electrophoresis. 2 μg of total RNA reverse transcribed using High-Capacity cDNA Reverse Transcription kit (Applied Biosystems, USA, Cat No. 4368814) according to the manufacturer’s instructions. The resulting cDNA was diluted ten times, and 2 μl was used as template for qPCR. qPCR reactions were set up in PowerUp^TM^ SYBR^TM^ Green PCR master mix (Thermo Scientific, PA, USA, Cat No. A25743) with gene specific primers (**Table 1**), and carried out in AriaMx Real-Time PCR System (Agilent, CA, US). The qPCR data generated in the form of Ct values were analysed as described in the figure legends.

**Table 1.**
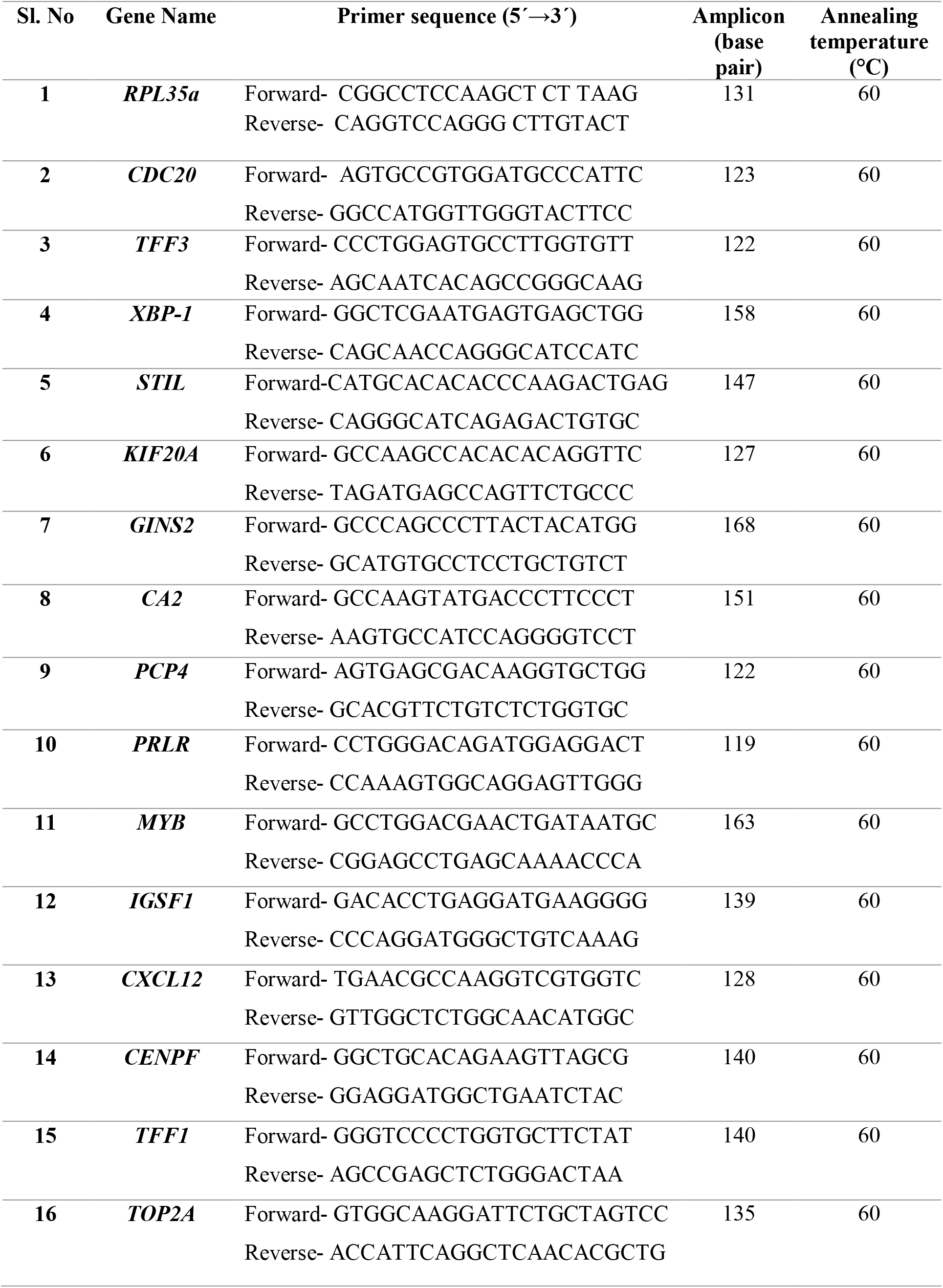
List of primers.

### 2.6. Chromatin immunoprecipitation

Chromatin immunoprecipitation was performed as described earlier (John Mary et al., 2020). 80 µg of chromatin sonicated in ChIP lysis buffer was precleared by incubating with Protein G plus-Agarose beads, which were pre-coated with bovine serum albumin (BSA) and herring sperm DNA. 5% of the pre-cleared chromatin samples were kept aside as input, and the remaining were incubated with ERα antibody (Cat. No. D8H8, Cell Signaling technology, MA, USA) or normal rabbit IgG antibody (BioBharati LifeScience, Kolkata, India) for 4 h, at 4°C. Immune complexes were pelleted by incubating with 20 μl of pre-coated protein G plus-Agarose beads for 2 h, at 4°C, followed by centrifugation. The pellets were washed extensively with a series of wash buffers. Immunoprecipitated chromatin samples were eluted and reverse cross-linked as described earlier (John Mary et al., 2020; Shivaswamy and Iyer, 2007). They were purified using Nucleospin Gel and PCR clean up kit from Machery-Nagel (Duren, Germany), and used in PCR reactions with primers capable of amplifying a 423 bp region containing an estrogen response element within the *TFF1* locus (Forward sequence: CATTGCCTCCTCTCTGCTCC. Reverse sequence: ACTGTTGTCACGGCCAAGCC).

### 2.7. RNA-seq

The materials and methods used for the RNA-seq experiment are described in detail in our earlier publication (Hatwik et al., 2023). A total of six biological replicate RNA samples isolated from MCF-7 cells treated with vehicle (0.1% DMSO, n = 3), or 10 μM EL (n = 3) were used for RNA-seq. The raw and processed data can be accessed through Gene Expression Omnibus (Accession number GSE216876). The read count data obtained from all the samples were analyzed by DESeq2 to identify differentially expressed genes.

### 2.8. Functional annotation and Gene Set Enrichment Analysis

Gene ontology (GO) was done via clusterProfiler package (Yu et al., 2012) in R, to identify the characteristic biological terms of RNA-seq data according to three functional categories; biological processes (BP), cell components (CC), and molecular function (MF). A separate GO analysis for up-regulated genes and down-regulated genes was performed. We employed GSEA software (Subramanian et al., 2005), with FDR correction of 25%, to find hallmark gene sets enriched in cells treated with 10 μM EL. Enrichment plots, and normalized enrichment scores (NES) were generated using additional R packages.

### 2.9. Statistical analysis

For identification of genes modulated by EL using DESeq2, we applied Wald statistic with α equal to 0.05, followed by FDR correction with a 5% cut-off. Viable cell count data generated in this study were analyzed by two-way ANOVA, only after confirming homogeneity of variances using the Levene test. Other multiple group data were analyzed by ANOVA. All statistical analyses were performed at 5% level of significance (p < 0.05). The results of all the statistical tests have been compiled in **Supplementary data 4**.

## 3. Results

### 3.1. Differential growth trajectories of MCF-7 and T47D cells cultured in M1 and M2 medium

A study on the effect of EL on MCF-7 or T47D cell-growth necessitated the choice of M2 as the treatment medium. M2, unlike M1, is devoid of phenol red, a known estrogenic agent (Welshons et al., 1988), and contains CS-FBS, which is depleted of steroid hormones. Prior to the study of the effect of EL, we examined the growth of these cell lines in M1 or M2 with respect to time. A total of four combinations were tested, namely MCF-7 in M1, MCF-7 in M2, T47D in M1 and T47D in M2. **Fig. 1** shows the growth trajectories for each of the four combinations. We applied two-way ANOVA to test the effect of time, combination of medium and cell-type, or their interaction. There were significant main effects of time (p ≈ 0), and combination (p ≈ 0). Notably, the interaction between time and combination was also significant (p ≈ 0), suggesting that the effect of time depended on the combination of medium and cell-type. A post-hoc test revealed that MCF-7 cells cultured for 120 h in M1 yielded significantly greater viable cell count compared to those cultured in M2 (p = 0; **Fig.1**, dark blue vs orange). In contrast, T47D cells outperformed MCF-7 cells in M1; the viable cell count of T47D cells after 120 h growth being significantly greater than MCF-7 cells (p = 0, **Fig. 1**, light blue vs dark blue). Furthermore, unlike MCF-7 cells, T47D cells appeared resilient in M2. This is reflected in the significantly higher viable cell count for T47D cells at the 120 h time-point, compared to MCF-7 cells (p = 0; **Fig. 1**; red vs orange).

**Figure 1.**
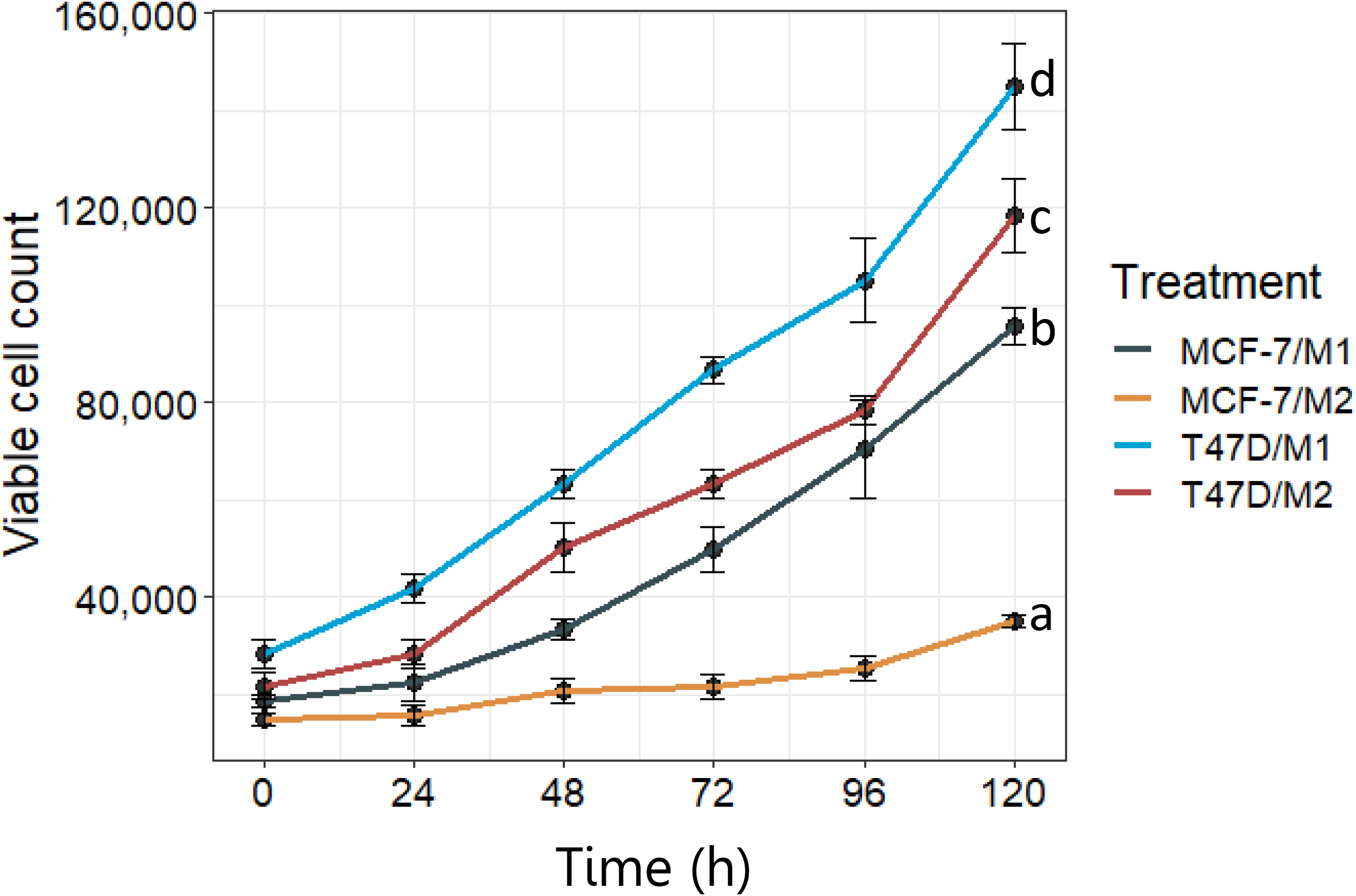
Growth trajectories of MCF-7 and T47D cells cultured in M1 or M2 medium. 7 × 10^3^ cells were seeded in 35 mm dishes using M1 medium. After 48 h, cells were cultured in M1 or M2. The culture dishes were replenished with fresh medium every 24 h. At the indicated time-points, total viable cell count in every dish was determined by the trypan-blue dye exclusion method as described in section **2.3**. The legend on the right shows the combination of cell-type and the medium in *cell-type/medium format*. The data were analyzed by two-way ANOVA to test for the effect of time, combination, or their interaction on total viable cell count followed by the Tukey’s HSD post-hoc test. The complete statistical analysis of the data is provided as **supplementary data 4**. Points on the graph represent mean total viable cell count (± SD, n = 3). a, b, c, and d, are letter codes showing statistical difference between combinations only for the 120 h data. The experiment was repeated two times with similar results.

### 3.2. 10 μM EL enhances the growth of MCF-7 and T47D cells

Dose-response experiments were performed to study the effect of EL on the growth of breast cancer cells in M2. ERα-positive (MCF-7, T47D), and ERα-negative (MDA-MB-231, MDA-MB-453) cells were treated with varying concentrations (0 to 40 μM) of EL, or 10 nM E2 in M2, for 0 (baseline viable cell count) or 120 h (end-point viable cell count). The results of the dose-response experiments are shown in **Fig. 2**. The viable cell-count data was analyzed by two-way ANOVA to test the effects of time, and treatment, or their interaction. As expected there was significant main effect of time (p ≈ 0) for all the cell lines; the total viable cell count after 120 h being significantly greater than the baseline at all concentrations of EL (**Fig. 2A-D**). There was significant main effect of treatment in all the cell lines (p = 0.036 for MDA-MB-231, and p ≈ 0 for others). Notably, the interaction between time and treatment was also significant in MCF-7, T47D, and MDA-MB-453 cells (p ≈ 0), but not in MDA-MB-231 cells (p = 0.08). In all the cell lines, there was no significant difference in the baseline viability determined at 0 h across all treatments (**Fig. 2A-D**; green dots). A post-hoc analysis showed that total viable count at 120 h, with 10 nM E2 was significantly greater than control, or any concentration of EL in MCF-7 and T47D cells (**Fig. 2A,B**; p << 0.001). There was no effect of 10 nM E2 on MDA-MB-231 or MDA-MB-453 cells (**Fig. 2C,D**). Notably, EL had differential effect on viable cell count over 120 h of treatment, depending on the concentration and the cell type. In MCF-7 cells, there was a significant increase in viable cell count with 10 and 40 μM EL (p ≈ 0, and p < 0.0001, respectively, **Fig. 2A**). In T47D cells, 10 μM EL significantly increased the viable cell count (p < 0.0001), while 40 μM EL significantly decreased the viable cell count (p < 0.05) with respect control (**Fig. 2B**). None of the concentrations of EL produced any significant increase or decrease in viable cell count in MDA-MB-231 cells (**Fig. 2C**), although in some replicate experiments, we did observe marginal increase or decrease in cell viability (*data not shown*). In MDA-MB-453 cells, 40 μM EL caused a significant decrease in viable cell count (p ≈ 0, **Fig. 2D**).

**Figure 2.**
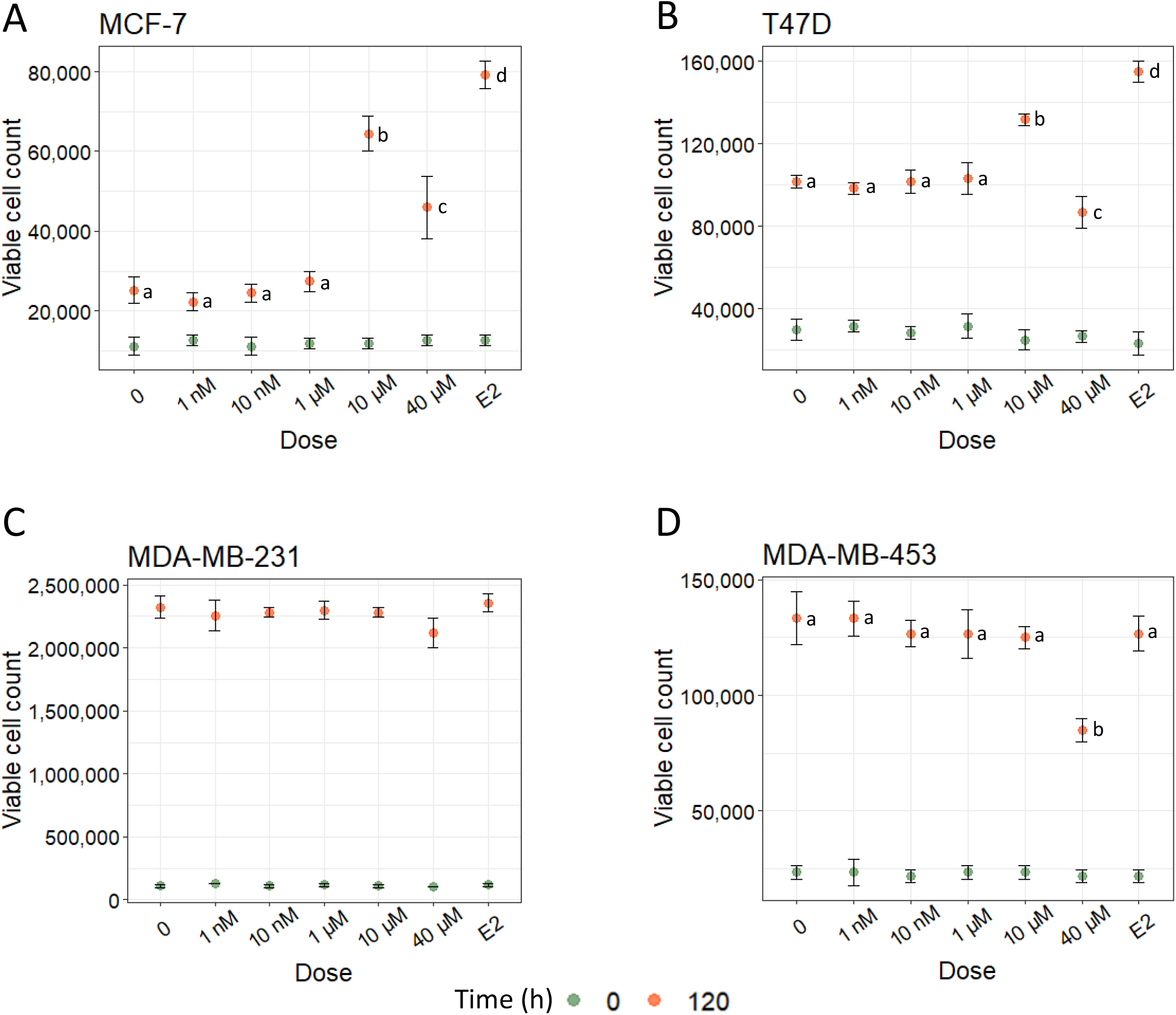
Dose-response of breast cancer cell lines to EL. 7 × 10^3^ cells were seeded in 35 mm dishes using M1. After 48 h, the cells were fed with M2, and maintained for 24 h. Thereafter, the cells were fed with M2 containing indicated concentrations (0-40 μM) of EL, or 10 nM E2, for a period of 0 (baseline viable count) or 120 h. The spent medium was replaced with fresh medium containing the treatments every 24 h. At the indicated time-points, total viable cell count in every dish was determined by trypan blue dye exclusion methods as described in section **2.3**. Each point represents mean viable cell count (± SD, n = 3). The data were analyzed by two-way ANOVA, to ascertain the main effects of time, treatments or concentration, or their interaction, followed by the Tukey’s HSD post-hoc test. The experiment was repeated two times with similar results.

We performed time-course experiments to capture the growth trajectories of breast cancer cell lines treated with vehicle, E2 or 10 μM EL in M2 medium. The data were analyzed by two-way ANOVA to examine the main effects of time, treatment or their interaction. Reflecting significant main effect of time, the viable cell count increased significantly in all the cell lines (p ≈ 0; **Fig. 3A,B**). The main effect of treatment was significant only in MCF-7 and T47D cells (p ≈ 0; **Fig. 3A,B**). Interestingly, the interaction between time and treatment in the two ERα-positive cell lines was significant (p ≈ 0, **Fig. 3A,B**) indicating difference in their growth trajectories in the presence of vehicle, E2 or EL. Differences in viable cell counts brought about by the treatments became magnified with time (**Fig. 3A,B**). Overall, at the 120 h time-point, EL produced significantly greater number of viable cells in MCF-7 and T47D cells (p < 0.001), albeit to a lesser extent than E2 (p ≈ 0). We examined the increase in viable cell counts obtained for both the cells lines after 120 h of vehicle or EL treatment. For MCF-7 cells the increase was 1.6 fold; the average viable cell count at 0, and 120 h being 9630 ± 1282 and 28148 ± 1283, respectively. In contrast, for T47D cells, the increase was 1.3 fold; the average viable count at 0, and 120 h being 26666 ± 2886 and 141666 ± 2886, respectively. Thus the growth response of MCF-7 cells to 10 μM EL appeared to be robust in comparison to that of T47D cells.

**Figure 3.**
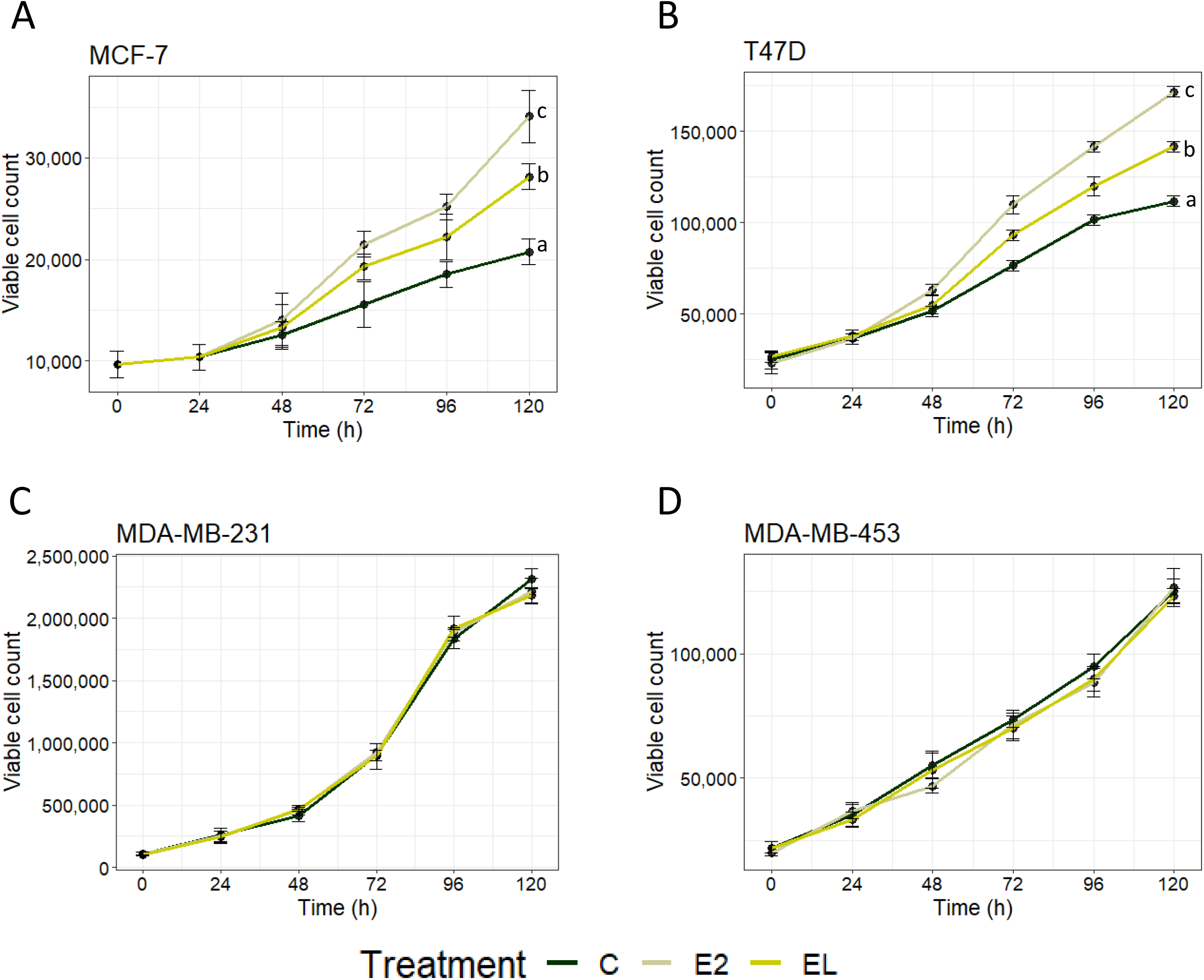
Growth trajectories of breast cancer cell lines treated with EL. 7 × 10^3^ cells were seeded in 35 mm dishes using M1. After 48 h, the cells were fed with M2, and maintained for 24 h. Thereafter, in dose-response experiments, the cells were treated with of 10 µM EL, or 10 nM E2 in M2, for the indicated periods of time. In time-course experiments, the cells were fed with M2 containing vehicle, 10 µM EL, or 10 nM E2 for the indicated periods of time. The spent medium was replaced with fresh medium containing the treatments every 24 h. At the indicated time-points, total viable cell count in every dish was determined by trypan blue dye exclusion methods as described in section **2.3**. Each point represents mean viable cell count (± SD, n = 3). The data were analyzed by two-way ANOVA, to ascertain the main effects of time, treatments or concentration, or their interaction, followed by the Tukey’s HSD post-hoc test. The experiment was repeated two times with similar results.

### 3.3. EL-mediated restoration of cell growth in MCF-7 cells is associated with maintenance of PCNA protein expression

Data presented in **Fig. 1** showed compromised growth of MCF-7, and the resilience of T47D cells in M2. We wanted to compare the growth of both cell lines grown in M2 alone, or M2 supplemented with 10 nM E2 or 10 μM EL, with those grown in M1. The data were analyzed by two-way ANOVA. As expected, the growth trajectory of MCF-7 with M2 medium had a gentle slope indicating compromised rate of cell proliferation (**Fig. 4A**, dark green). The viable cell count at 120 h time-point for cells grown in M2 medium supplemented with E2 was comparable to that in M1 (**Fig. 4A**, grey vs red). EL treatment in M2 significantly increased the viable cell count of MCF-7 cells (p ≈ 0, **Fig. 4A**, dark green vs yellow), albeit to a lesser extent compared to that achieved with M1 (p = 0; **Fig 4A**, red vs yellow), or M2 supplemented with E2 (p ≈ 0; **Fig. 4A**, grey vs yellow). Since, the growth of T47D was already robust in M2 (**Fig. 4B**, dark green), the impact of EL on its growth in M2 was not as robust as seen in MCF-7 cells (p = 0.004; **Fig 4B**, dark green vs yellow). Overall, viable cell counts for the 120 h time-point for T47D cells grown in M1, and M2 supplemented with EL or E2 were significantly different (p = 0.004 and p= 0.001; **Fig 4B**, grey vs yellow, and red vs yellow, respectively). The growth kinetic data correlated with the levels of PCNA protein expression in both the cell lines. Western blot analysis showed that after 72 h of culture in M2, PCNA protein level in MCF-7 cells falls (**Fig. 4C**, lanes 5 and 7), but remains stable when cultured in M1 medium (**Fig. 4C**, lanes 1 and 3). Furthermore, when cultured in M1 medium for 72 h, 10 μM EL had no effect on PCNA levels (**Fig. 4C**, lanes 1 and 4). However, in the presence of M2 medium, 10 μM EL appeared to restore PCNA protein expression (**Fig 4C**, lanes 5 and 8). In contrast, irrespective of the medium (M1 or M2), EL did not have any impact on PCNA protein expression in T47D cells (**Fig. 4D**).

**Figure 4.**
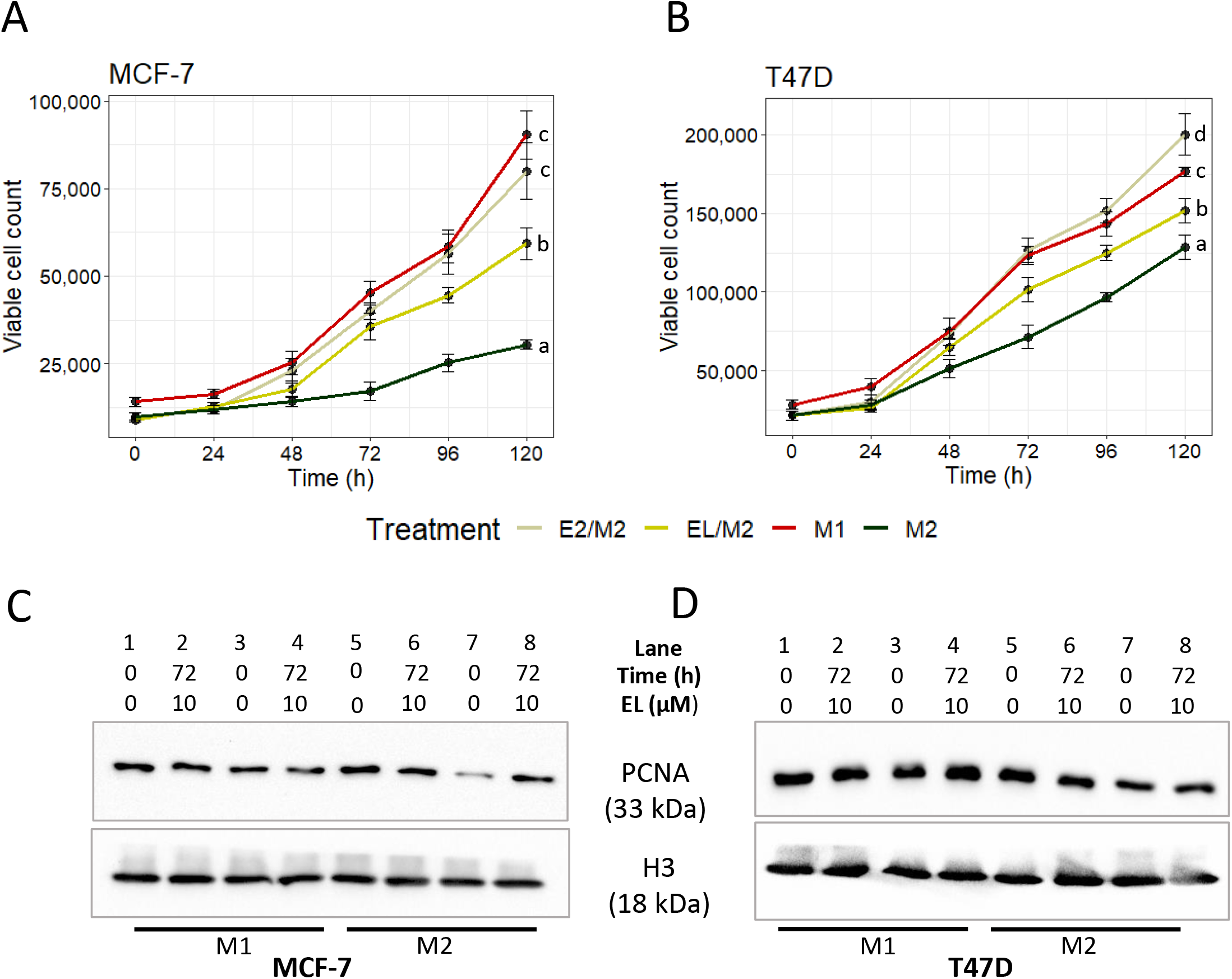
10 μM EL restores PCNA expression associated with MCF-7 cell growth in M2 medium. A, B. Growth trajectories of MCF-7 (A) and T47D (B) cells. 7 × 10^3^ cells were seeded in 35 mm dishes using M1. After 48 h, the cells were fed with M2, and maintained for 24 h. Thereafter the cells were grown in M2 containing vehicle (0.1% DMSO), 10 μM EL, or 10 nM E2 for indicated periods of time. The growth trajectories were compared to the cells maintained in M1 medium. At the indicated time-points, total viable cell count in every dish was determined by trypan blue dye exclusion methods as described in section 2.3. Each point represents mean viable cell count (± SD, n = 3). The data were analyzed by two-way ANOVA, to ascertain the main effects of time, treatments or concentration, or their interaction, followed by the TukeyHSD post-hoc test. C, D. Western blot analysis of PCNA protein expression. MCF-7 (C) and T47D (D) cells were treated with vehicle (0.1% DMSO), or 10 μM EL in M1 or M2 for 0 or 72 h. Total protein was extracted and 30 μg of each sample was subjected to western blot analysis using PCNA-specific antibody, as described in section **2.4**. H3 served as an internal control.

### 3.4. Tamoxifen blocks EL-mediated increase in viable cell count in MCF-7 and T47D cells

We studied the effect of tamoxifen (Tam) on EL induced growth of MCF-7 and T47D cells. Cells were treated with vehicle or 10 μM EL, alone or in combination with 1 μM tamoxifen for 0 (baseline) or 120 h. The data were analyzed by two-way ANOVA to test the main effects of time, treatment, or their interaction. There were significant main effects of time, treatment and interaction in both the cell lines. The baseline viability of MCF-7 or T47D cells in the treatment groups was not significantly different (red dots, **Fig. 4A,B**) The significant effect of time is reflected in the significant increase in viable cell count in all treatment groups (**Fig. 4A,B**). The post-hoc analysis revealed that after 120 h of treatment, the viable cell count was significantly greater with 10 μM EL compared to control (**Fig. 5A,B**; C vs EL). While 1 μM tamoxifen alone had a significant negative effect on the viable cell count (**Fig. 5A,B**; C vs TAM), it significantly reduced the viable cell count brought about by EL (**Fig. 5A,B**; EL vs EL+TAM).

**Figure 5.**
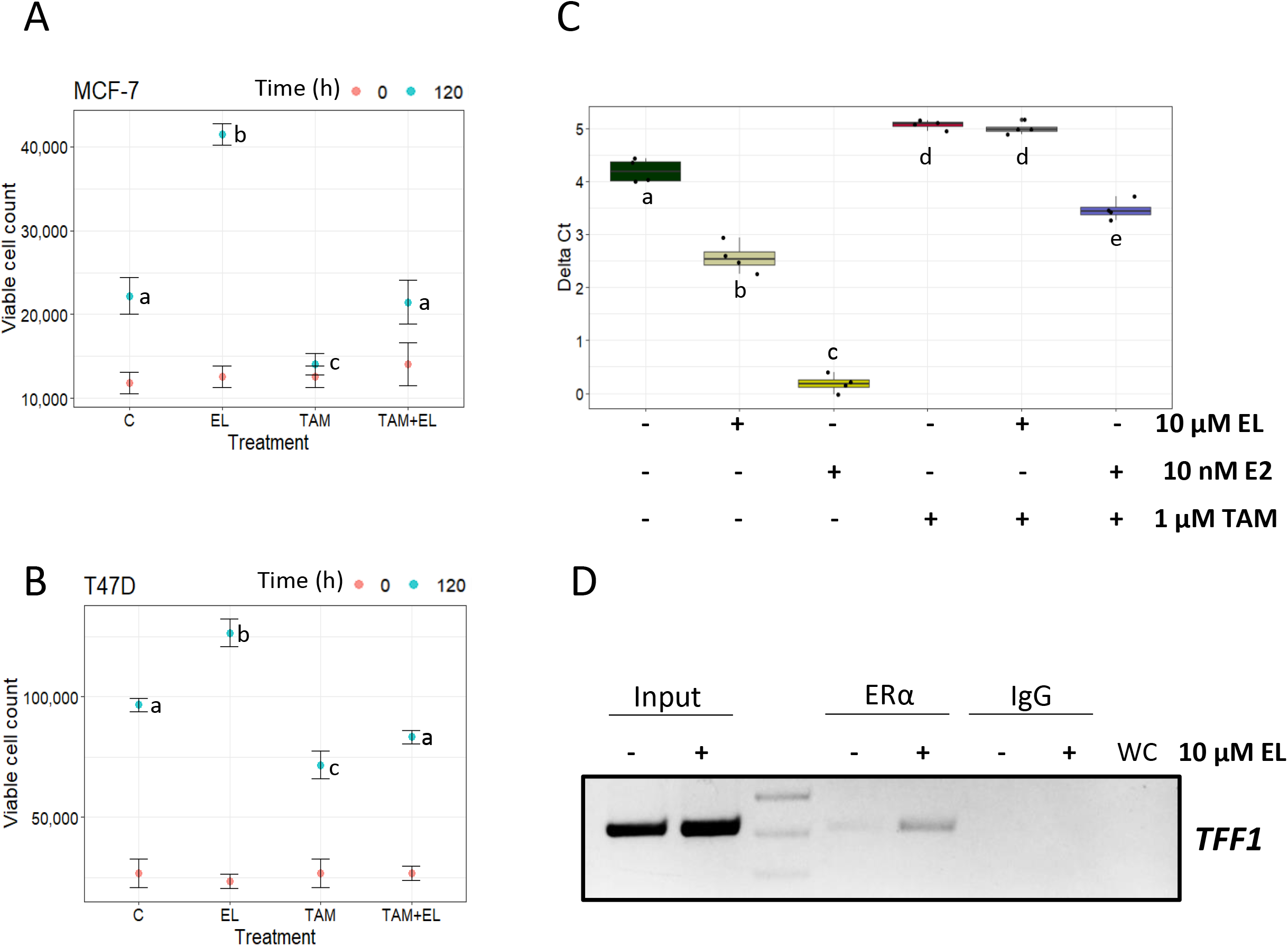
Effect of Tamoxifen on 10 μM EL-mediated increase in cell viability and *TFF-1* mRNA. A, B. 7 × 10^3^ MCF-7 (A) or T47D (B) cells were seeded in 35 mm dishes using M1. After 48 h, the cells were fed with M2, and maintained for 24 h. The cells were then fed with 10 μM EL, 1 μM tamoxifen, or both for 0 (baseline viability) or 120 h. Control cells were treated with vehicle (0.1% DMSO). The data were analyzed by two-way ANOVA to test the effects of time, Treatment or their interaction. TukeyHSD was applied for the post-hoc test for ascertaining significant difference between groups. a, b, c, and d, are letter codes showing statistical difference between pairs of treatments only for the 120 h data. C. Total RNA was isolated from MCF-7 cells treated with vehicle, 10 μM EL, 1 μM tamoxifen, or both for 72 h and subjected to RT-qPCR analysis. For each sample the average Ct value for a *TFF1* mRNA (Ct^TFF1^), and *RPL35a* (Ct^RPL35a^) obtained from four reactions were determined. The difference Ct^TFF1^ - Ct^RPL35a^, which is referred to as ΔCt was considered as a measure of the normalized variant expression; serving as an internal control. Thus, higher the ΔCt value, lower the expression in a given sample. The data were analyzed by ANOVA followed by TukeyHSD. Bars represent mean relative expression (± SD, n = 3). D. Sonicated chromatin samples prepared from vehicle- or 10 μM EL-treated cells were immunoprecipitated with non-specific IgG or ERα-specific antibody. Immunoprecipitated DNA samples were subjected to PCR using primer pairs designed to specifically amplify the region encompassing an estrogen response element.

### 3.5. 10 μM EL enhances the expression of TFF-1 mRNA in MCF-7 cells

The mechanism of estrogen-mediated induction of *TFF1* mRNA is well known (Berry et al., 1989). It involves ERα, and its binding to the estrogen response element in the *TFF-1* locus. Tamoxifen blocks estrogen mediated induction of *TFF-1* mRNA (**Fig. 5C**; boxes 1, 3 and 6). 10 μM EL also increased the *TFF1* mRNA levels in MCF-7 (**Fig. 5C**; boxes 1 and 2) cells, although the fold-increase was significantly lesser compared to that brought about by 10 nM E2 (**Fig. 5C**; bars 2 and 3). Tamoxifen blocked the EL-mediated induction of *TFF1* mRNA (**Fig. 5C**; boxes 2 and 5). Furthermore, EL-mediated induction of *TFF-1* RNA was associated with increased binding of ERα to the estrogen response element within the *TFF-1* locus (**Fig. 5D**).

### 3.6. Impact of EL on MCF-7 transcriptome

The aforementioned results highlighted the proliferation inducing effect of 10 μM EL on ERα-positive breast cancer cells. To capture the transcriptomic response associated with EL-induced proliferation, we employed the next generation sequencing technology (RNA-seq). MCF-7 cells were treated with vehicle (0.1% DMSO) or 10 μM EL in M2 medium, and the total RNA samples were subjected to RNA-seq on the Illumina platform as described earlier (Hatwik et al., 2023). The read count data, which was obtained after adaptor trimming and filtering, followed by read alignment, was analyzed to determine differentially expressed genes using DESeq2 package in R. As shown in the volcano plot presented in **Fig. 6A**, 1141 genes were modulated by 10 μM EL in MCF-7 cells, based on a threshold value of 0 for log_2_FC, and 0.05 for Wald statistic and FDR (**Supplementary data 1**). These comprised 727 upregulated and 414 downregulated genes as illustrated in the heatmap in **Fig 6B**. The top 25 upregulated, and 25 downregulated genes are presented in **Table 2** and **Table 3**, respectively.

**Figure 6.**
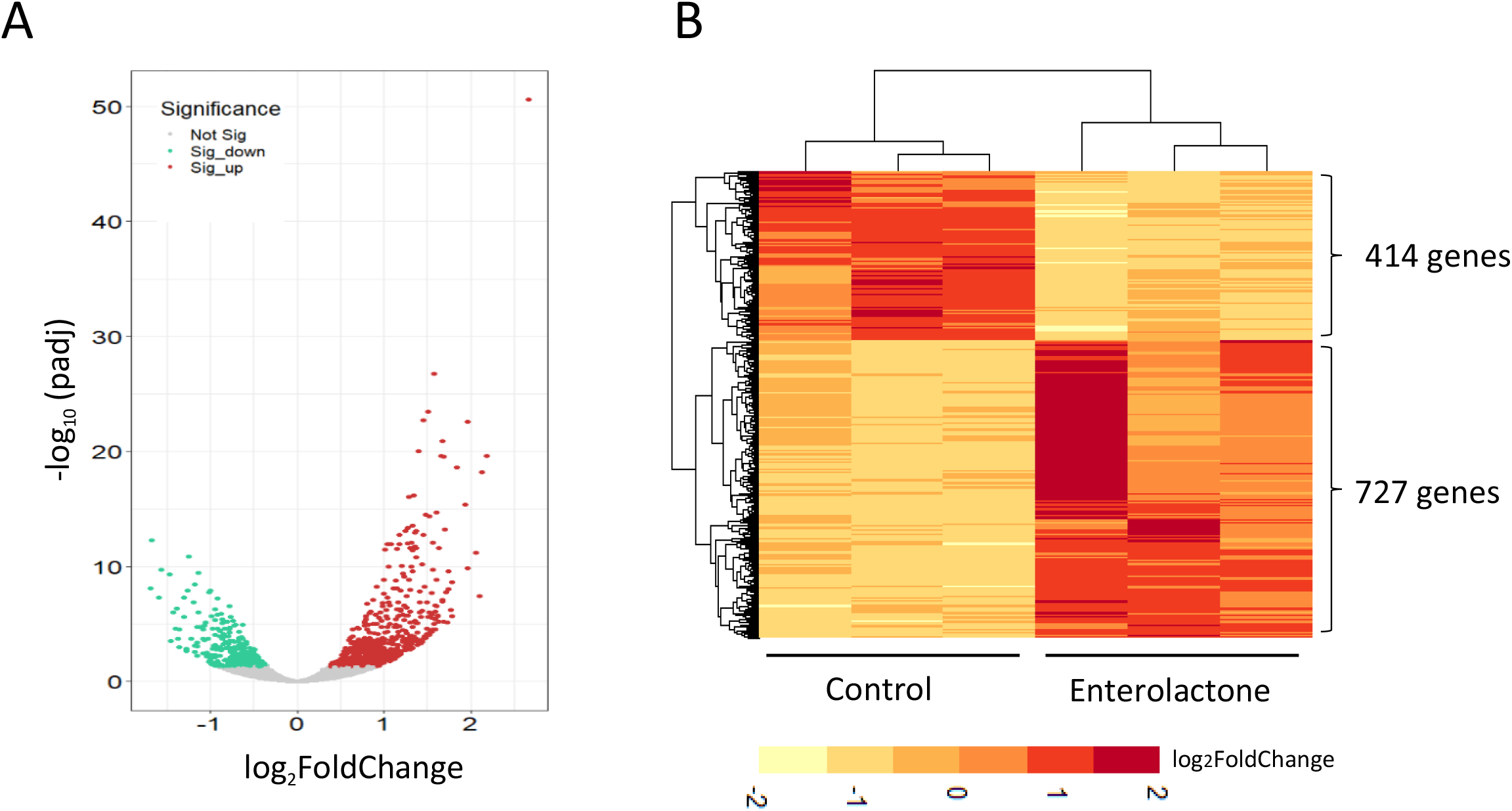
Summary of RNA-seq data. (A) Volcano plot. Genes significantly regulated by 10 µM EL are represented as colored dots according to their –log_10_P_adj_ and log_2_FC. Red dots (n = 727) represent the upregulated genes, while the green dots (n = 414) represent the downregulated genes. (B) Expression heatmap of significantly regulated genes by EL. The upregulated genes (log_2_FC>0) are represented in shades of red, downregulated (log_2_FC<0) are represented in shades of yellow.

**Table 2.**
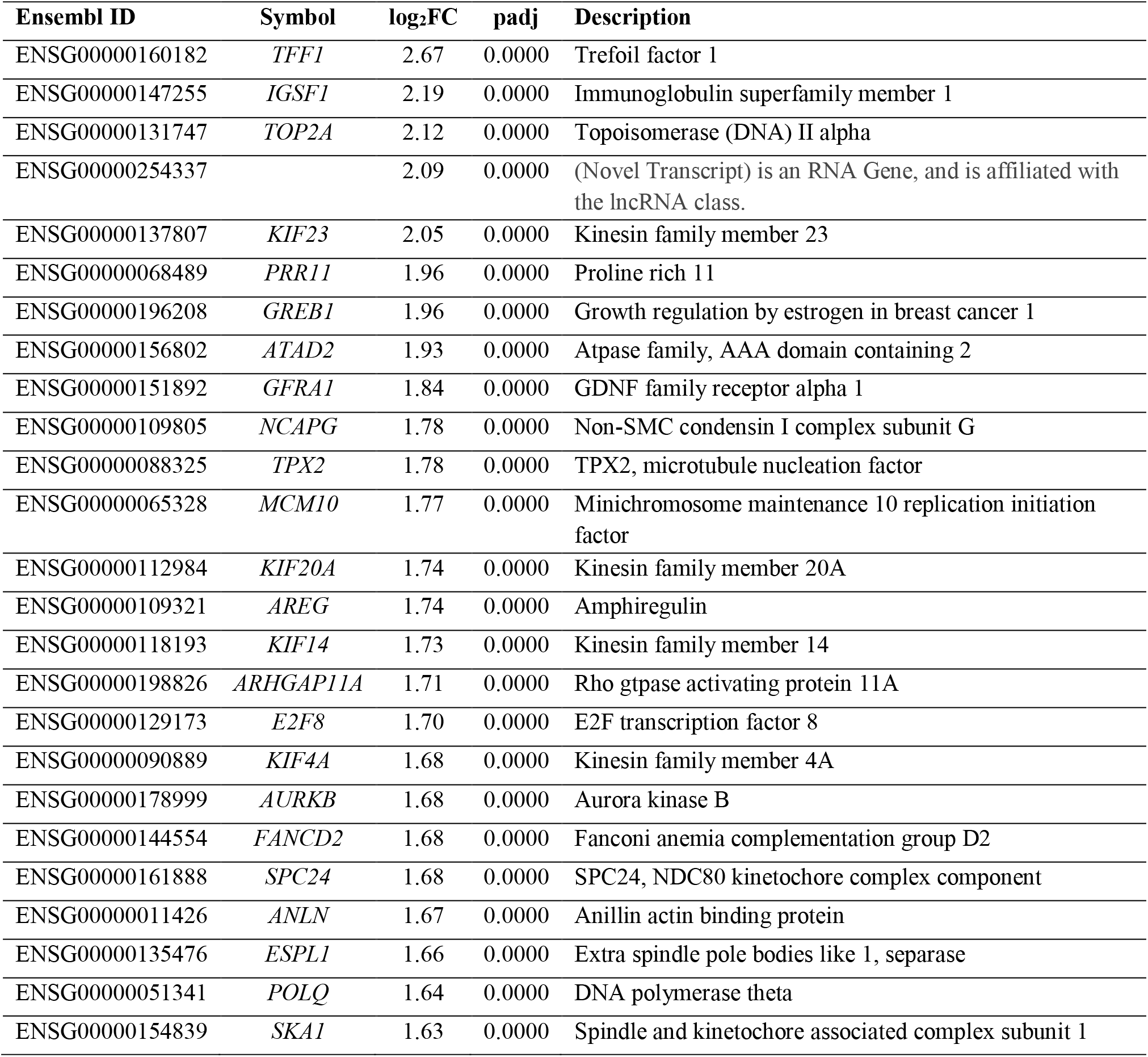
List of top 25 upregulated genes upon 10 µM EL treatment for 72 h in MCF-7 cells.

**Table 3.**
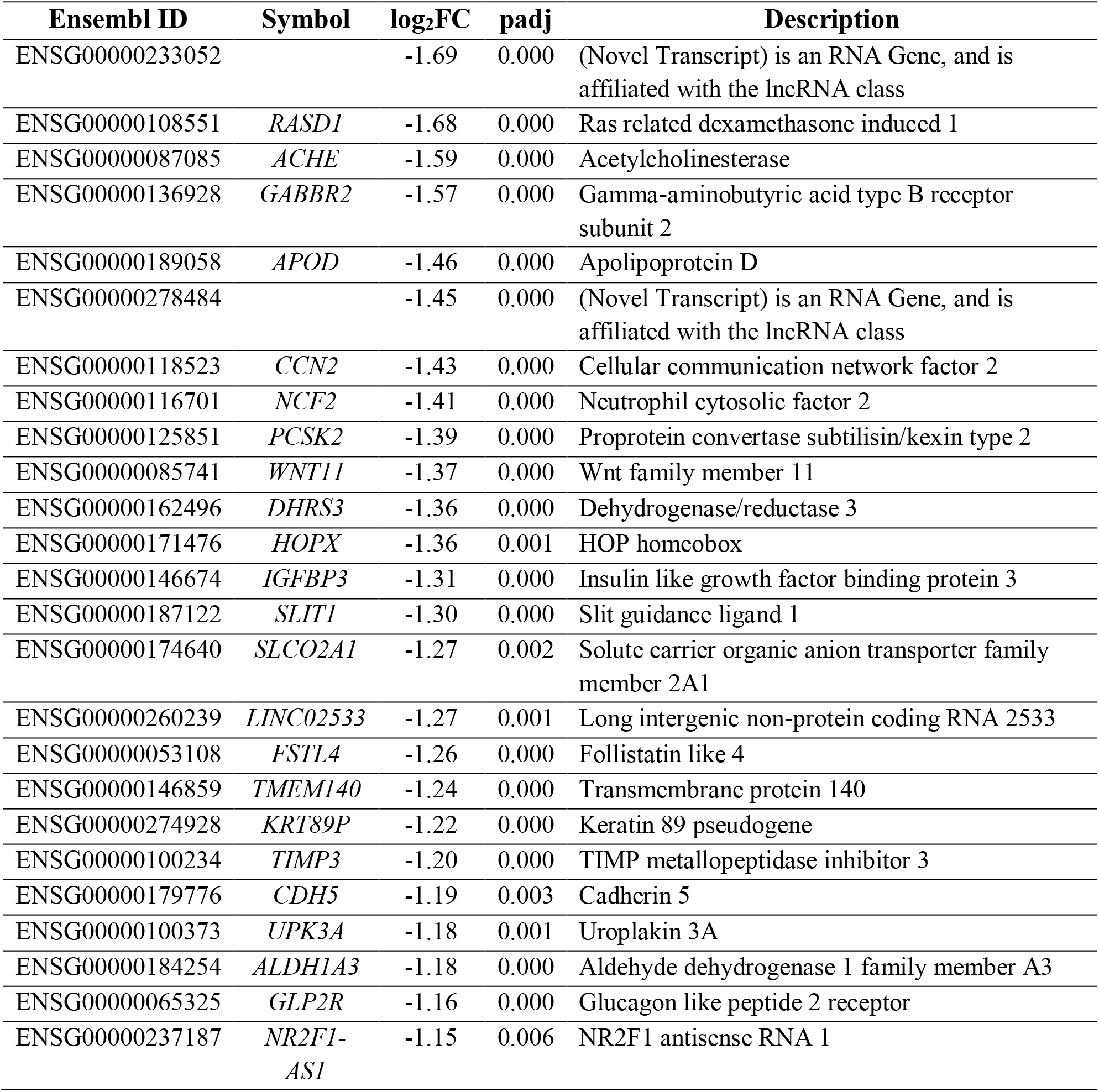
List of top 25 downregulated genes upon 10 µM EL treatment for 72 h in MCF-7 cell.

### 3.7. Gene Ontology analysis

On the derived set of significantly modulated genes, we ran the GO analysis. This enabled identification of specific groups represented in EL-modulated genes, which fell within the broader categories of cellular component, biological processes, and molecular function. The results of the analysis are presented as **Supplementary data 2**. The top GO-terms over-represented in the EL-induced, or -repressed genes are shown in **Fig. 7A**. Under the category of molecular function, over-represented terms associated with the EL-induced genes included *microtubule motor activity*, *ATP dependent activity*, *DNA binding*, and *tubulin binding,* and *microtubule binding* (**Fig. 7A**, left panel, grey bars). On the other hand, *calcium ion binding* was over-represented in those associated with EL-repressed genes (**Fig. 7A**, right panel, grey bar). Under the category of cellular compartment, over-represented terms associated with EL-induced genes included *chromosome*, *chromosomal region*, *centromeric region*, and *condensed chromosome* (**Fig. 7A**, left panel, green bars). In contrast, those associated with EL-repressed genes included *collagen containing extracellular matrix*, *extracellular matrix*, *external encapsulating structure*, *extracellular region*, and *cell periphery* (**Fig. 7A**, right panel, green bars). Over-represented terms under the broad category of biological processes associated with EL-induced genes included *chromosome segregation, cell division*, *cell cycle process*, *mitotic cell cycle process,* and *cell cycle* (**Fig. 7A**, left panel, yellow bars). In contrast, those associated with EL-repressed genes included *regulation of signalling, regulation of cell communication, regulation of signal transduction, blood vessel morphogenesis, and regulation of multicellular organismal process* (**Fig. 7A**, right panel, yellow bars).

**Figure 7.**
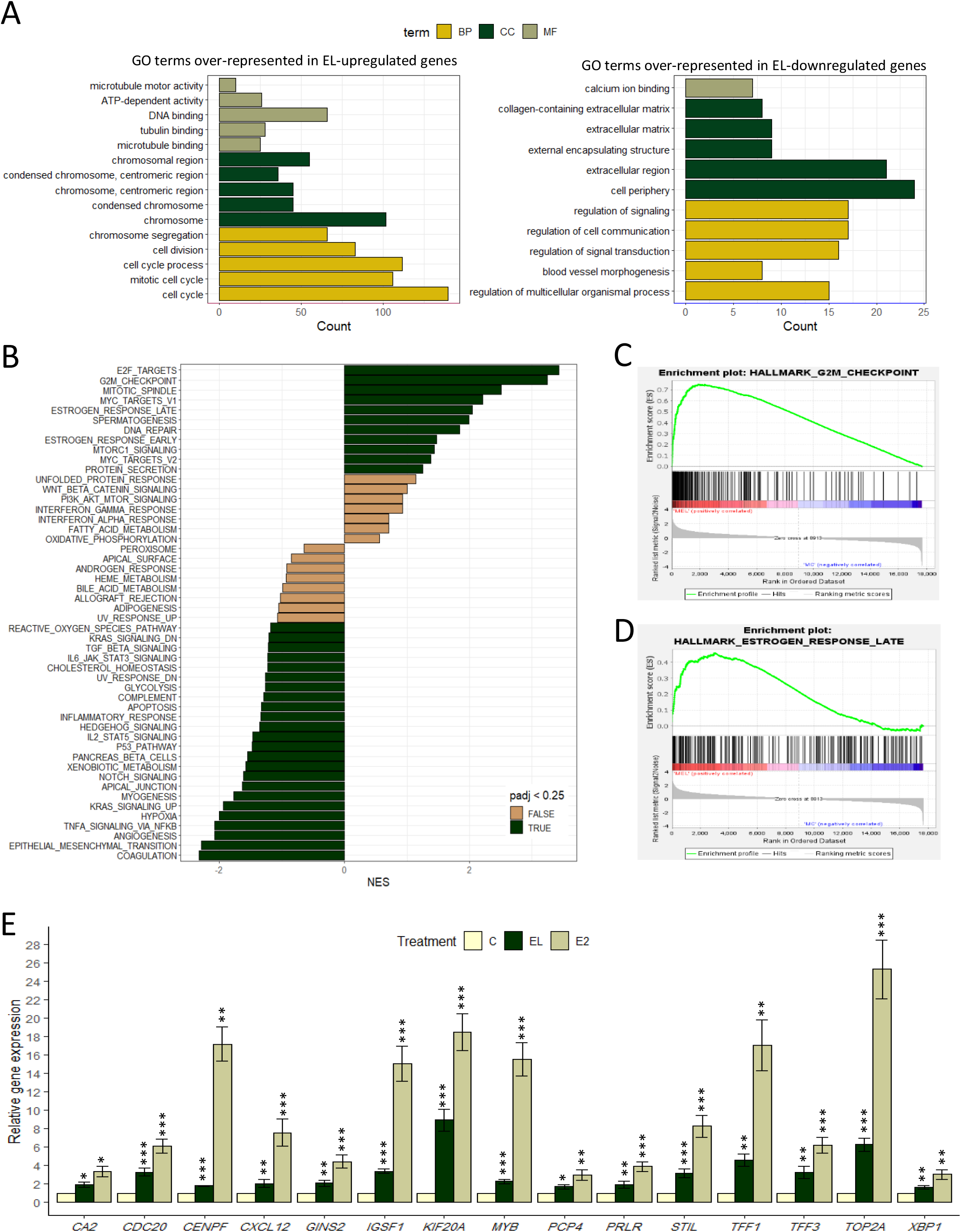
Enrichment analysis of EL-modulated genes in MCF-7. Top enriched GO terms of upregulated (A, left panel) and downregulated (A, right panel) EL-differentially expressed genes in biological processes (BP), cellular components (CC), and molecular functions (MF). clusterProfiler package in R was assigned for the analysis with FDR of 5%. (B) Gene set enrichment analysis. The analysis was performed using GSEA software. Green bars represent the hallmark gene sets enriched by EL based on 25% FDR cut-off. Respectively, (C) and (D) represent enrichment plots of G2/M checkpoints and estrogen response late pathways. (E) Validation of selected leading edge genes in G2/M checkpoint and estrogen-response-late gene-sets. Total RNA from vehicle-, 10 μM EL-, or 10 nM E2-treated samples were subjected RT-qPCR analysis. *RPL35a* served as internal control. The expression in EL- or E2-treated samples was expressed relative to those in vehicle controls using the ΔΔCt method (Livak and Schmittgen, 2001). Bars represent mean relative gene expression (± SD). For each gene, the fold induction by EL or E2 were analyzed by t-test. *p < 0.05, ** p < 0.01, ***p < 0.001.

### 3.8. Gene Set Enrichment Analysis (GSEA)

We applied GSEA to identify hallmark gene sets which were enriched in EL-regulated genes. The overall output of GSEA is presented in **Supplementary data 3**. The significantly enriched gene-sets, based on FDR < 25%, are indicated by green bars in **Fig. 7B**. The enrichment of G2/M checkpoint genes is noteworthy, which correlates with the ability of 10 µM EL to induce growth of MCF-7 cells. The enrichment plot for the G2/M checkpoint gene-set is shown in **Fig 7C**. There were 115 leading edge genes within this gene-set (**Supplementary data 3**). Out of these, the modulation of *CDC20*, *CENPF*, *TOP2A*, *STIL*, and *GINS2* were validated by RT-qPCR (**Fig. 7E**). The EL-regulated genes were also enriched in the estrogen-response-late, estrogen-response-early, and DNA repair gene sets. The enrichment plot for the estrogen-response-late gene set is shown in **Fig. 7D**. There were 69 leading edge genes in this set (**Supplementary data 3**). Out of these, we validated the modulation of *TFF1*, *TFF3*, *CA2*, *CDC20*, *CXCL12*, *GINS2*, *IGSF1*, *KIF20A*, *MYB*, *PCP4*, *PRLR*, *STIL*, *TOP2A*, and *XBP1* by RT-qPCR (**Fig. 7E**). We also examined the effect of 10 nM E2 on the expression of the EL-modulated genes. In general, the induction brought about by 10 μM EL was lower than that produced by 10 nM E2 (**Fig. 7E**).

## 4. Discussion

In the light of the epidemiological data on the possible link between enterolactone exposure and breast cancer risk (Buck et al., 2010), investigations into the pro- or anti-estrogenic effects of EL (Mali et al., 2019; Zhu et al., 2017) cannot be overlooked. Pro-estrogenic molecules are expected to induce proliferation, and modulate the expression of estrogen target genes in ERα-positive and estrogen responsive cells. EL qualifies as a pro-estrogenic molecule on both counts. Studies on ERα-positive breast cancer cells have repeatedly demonstrated proliferative effect of EL only upto 10 μM, but not higher concentrations (Mousavi and Adlercreutz, 1992; Sathyamoorthy et al., 1994; Welshons et al., 1987). The present study confirms this result via elaborate dose-response (Fig. 2) and time-course experiments (Fig. 3). Zhu et al. (2017) have demonstrated that EL-mediated proliferation of MCF-7 cells is associated with activation of cyclin-CDK complexes (Zhu et al., 2017), which is typical of estrogen-mediated induction of cell proliferation (Foster et al., 2001). Our analysis of PCNA expression, a marker of cell proliferation, provides an independent perspective on EL action. The time-dependent fall in PCNA expression in MCF-7 cells cultured in M2, a medium depleted of growth supporting hormones, is restored by 10 μM EL. This supports the growth promoting effect of 10 μM EL in MCF-7 cells, although the effect is significantly lesser comparable to 10 nM E2. Notably, T47D cells, which inherently proliferate faster than MCF-7 cells (**Fig 1**), were resilient to M2 medium and did not show any fall in PCNA levels (**Fig 4D**). This could be due to autocrine proliferative mechanisms that continue to support the growth of T47D cells, despite hormone depletion. As a result, the growth response of MCF-7 cells to 10 μM EL is more robust compared to that of T47D cells (**Fig2 A, and 3A**), which justifies the choice of MCF-7 cells as a model to capture gene expression modulation associated with EL-induced cell proliferation.

Physical binding of EL to ERα (Penttinen et al., 2007) raises the possibility that its actions are mediated, at least in part, via the classical estrogen receptor in ERα-positive breast cancer cells. It leads to the hypothesis that 10 μM EL-mediated induction in MCF-7 or T47D proliferation could be mediated via ERα. Our results show that 10 μM EL does not enhance the proliferation of the ERα-negative MDA-MB-231 or MDA-MB-453 cells (Fig. 2 and 3), which strengthens the hypothesis. Zhu et al. (2017) showed that ICI182780 (fulvestrant), which selectively degrades ERα, inhibits the proliferation of MCF-7 cells induced by 10 μM EL. Our observation that tamoxifen, a selective estrogen receptor modulator, prevents enhanced proliferation of MCF-7 by 10 μM EL, complements their result. EL induces ERα transcriptional activity in a dose-dependent manner (Carreau et al., 2008). It induces Trefoil factor-1 (Sathyamoorthy et al., 1994), and PR expression in MCF-7 cells, and induces prolactin synthesis in pituitary cells (Welshons et al., 1987). *In vivo*, it produces uterine stromal edema, one of the signs of estrogenic activity (Penttinen et al., 2007). Notably, these effects are mediated by EL at a concentration of 10 μM. Furthermore, antiestrogen inhibits EL-mediated induction of progesterone receptor (PR) in MCF-7 cells (Welshons et al., 1987). Here we show that 10 μM EL significantly induces the expression of Trefoil factor 1 in MCF-7 cells via a mechanism that involves ERα binding to the estrogen response element within the *TFF1* locus (Fig. 5). Through a microarray based expression profiling of a limited set of estrogen responsive genes, Zhu et al. (2017) showed a significant correlation between the effects of E2 and EL on gene expression. Taken together these are compelling evidences in favour of a significant role for ERα in EL-induced cell proliferation and alteration of gene expression at 10 μM concentration.

The present study, while presenting the transcriptomic changes associated with EL-mediated proliferation, addresses the lacuna regarding the genome-wide effects of EL. The functional diversity in the genes that are modulated by EL is apparent from the results of GO, and GSEA. Induced proliferation entails rapid cell cycle progression and higher frequency of cell division. In the face of induced proliferation, cells channelize energy for DNA synthesis, chromosomal duplication and segregation, and negotiation of cell cycle checkpoints. Cellular processes that entail communication with the environment are not a priority. Over-representation of terms, such as such as those associated with chromosomes and cell cycle (Fig. 7A, left panel) in the EL-induced genes, and over-representation of terms associated with membrane, and signalling (Fig. 7A, right panel) in the EL-downregulated genes essentially reflect this notion.

The enrichment of the hallmark G2/M checkpoint genes on the one hand, captures the systemic shift towards proliferative state under the influence of 10 μM EL. On the other hand, enrichment of estrogen-response-late and estrogen-response-early genes reflect the involvement of the ERα signalling axis. It is worth noting that *CDC20*, *STIL, TOP2A*, and *GINS2* were among the leading edge genes in both sets. CDC20, cell cycle modulator, controls the chromosomal segregation during mitosis (Hartwell et al., 1973). It is over-expressed in breast tumors compared to normal tissues (Jiang et al., 2011), and known as an oncoprotein (Karra et al., 2014). GINS2 is a subunit of one of the core components of eukaryotic replicative helicase, whose role is involved in DNA replication by opening the double-stranded DNA at replication fork (Onesti and MacNeill, 2013). GINS2’s expression is higher in breast cancer tissues compared to normal ones (Yu et al., 2020), and it was identified as metastasis promoting protein in breast cancer patients (Thomassen et al., 2009). Further, TOP2A, is a nuclear enzyme involved in DNA replication and transcription via controlling DNA topology (Wang, 2002). High TOP2A in breast cancer patients is thought to negatively affect caner prognosis (Zaczek et al., 2012), such as increased tumor grade and Ki67 index (An et al., 2018). Besides, Centriolar Assembly Protein (STIL), is a cytosolic protein, promotes centriole duplication in proliferating cells (Vulprecht et al., 2012). It is upregulated in many tumors including breast tumors (Li et al., 2022). The induction in the expression of the aforementioned genes suggests estrogen like effect of EL at 10 μM concentration. It also aligns with the intertwining of genomic and proliferative effects of estrogen. Both viability assays, and gene expression studies have shown repeatedly, that the effect produced by EL is significantly lower than that produced by E2. There is a substantial difference in their affinities to the ERα (Mueller et al., 2004), which possibly underlies the quantitative difference in their effects. However, being different chemical entities, they may induce conformational changes in ERα, differently, though subtly, just like in case of selective estrogen receptor modulators. That may lead to differential effects on gene expression. This begs an independent interrogation into the extent of overlap between genes regulated by EL and E2, both qualitatively and quantitatively.

Thus, the present study provides a genomic perspective on the estrogenic effect of EL in MCF-7 cells. It does not address the opposite effect of EL on breast cancer cell proliferation at higher concentrations, which is also important in understanding the epidemiological difference in the effects of EL in pre- and post-menopausal women. However, the results of this study bring to the fore a caveat regarding the breast cancer-preventing effect of EL. It is likely that the true effect of EL on the mammary gland is a function of the concentration EL, and the existence of a functional estrogen-ERα signalling axis.

## Acknowledgment

The work was supported by financial assistance from Science and Engineering Research Board, Department of Science and Technology, Govt. of India (File No. CRG/2020/002109). Infrastructural support from Department of Biosciences and Bioengineering, IIT Guwahati is acknowledged. JH acknowledges the research fellowship from Al-Baath University, Ministry of Higher Education and Scientific Research, Syrian Arab Republic (Grant no. /188/B, Date: 20.06.2017).

